# Genome-resolved biogeography reveals multidimensional structuring of freshwater giant viruses across global deep lakes

**DOI:** 10.64898/2026.08.06.743156

**Authors:** Liwen Zhang, Michaela M Salcher, Morimaru Kida, Hideo Oyagi, Yoshikuni Hodoki, Atsushi Toyoda, Ken Kurokawa, Hideyuki Tamaki, Shin-ichi Nakano, Hiroyuki Ogata, Yusuke Okazaki

## Abstract

Giant viruses (GV) are increasingly recognized as important ecosystem regulators. While metagenomics has uncovered extensive GV diversity, the global distributions of individual species and the biogeographic processes driving the pattern remain poorly understood. Here, we reconstructed GV metagenome-assembled genomes (MAGs) from 35 globally distributed deep freshwater lakes spanning five continents, aiming to identify their biogeographic patterns. The resulting 1663 non-redundant MAGs significantly expanded the known freshwater GV diversity, with ∼84% lacking a previously reported species representative. These MAGs were grouped into cosmopolitan and geographically restricted lineages. We identified 27 cosmopolitan GV species spanning multiple viral lineages, including families of *Imitervirales*, *Pimascovirales*, and mirusviruses order *Styxvirales*. The cosmopolitan species were characterized by their larger genomes and expanded gene repertoires of host-interaction functions, which may facilitate interactions with diverse hosts and contribute to their global distributions. The presence of geographically restricted species and the stronger distance-decay in community similarity observed in freshwater than marine ecosystems suggest that physical connectivity between ecosystems is an important factor influencing GV dispersal. We identified 312 and 177 GV MAGs almost exclusively associated with the epilimnion and hypolimnion, respectively. This water-layer preference of individual MAGs was highly consistent across lakes, suggesting conserved vertical partitioning in association with the thermal stratification of the water column. Overall, our findings reveal that GV biogeography in deep freshwater lakes is structured by the combined influence of horizontal dispersal limitation, vertical partitioning, and lineage-specific evolutionary histories.

## Introduction

Giant viruses (GVs) belonging to the phyla *Nucleocytoviricota* and *Mirusviricota* constitute phylogenetically diverse and ecologically important eukaryotic double-stranded DNA viruses widely distributed across various ecosystems [1–6]. However, our understanding of GV biogeography remains limited. In aquatic systems, biogeographic patterns may emerge from the interaction of dispersal and colonization processes, vertical partitioning, and temporal dynamics, together defining a multidimensional framework of microbial biogeography [7,8]. Although these processes have been extensively described for bacteria, archaea, and microbial eukaryotes [8–12], those of GVs remain poorly understood. Early studies based on hallmark genes (e.g. *polB* and *MCP*) of marine GVs revealed their global distributions [1,13]. Later, metagenomic genome-resolved analyses improved the phylogenetic resolution and uncovered strong biogeographic structuring of GV communities along latitudinal and thermal gradients, including a pronounced distinction between polar and non-polar communities and repeated evolutionary transitions into polar-adapted environments [3,14]. Other studies further revealed that water layers also represent an important axis of GV biogeography, with identification of GVs unique to the dark water below 150 m in four oceanic transects [15] and exclusively abundant in the deep layers of the North Pacific Subtropical Gyre [16]. Beyond genomic surveys, transcriptomic analyses have revealed distinct GV lineages that are actively infecting hosts in aphotic deep-sea environments by modulating their gene expression pattern to thrive in these harsh habitats [17].

In contrast to the ocean, GV biogeography in freshwater ecosystems remains poorly characterized. Most existing studies have focused on individual lakes [5,18] or regional datasets [19], limiting our ability to determine whether and how freshwater GV communities are structured across water layers and geographic regions. Regarding vertical structuring, a recent study revealed that GV communities were strongly structured along water layers in a deep lake, with distinct GV lineages exclusively associated with either the epilimnion or hypolimnion [5]. At the geographic scale, a global investigation on a cryptophyte-infecting imitervirus species has revealed its cosmopolitan distribution across multiple freshwater systems [20]. Occasional observation of almost identical GV genomes across continents [5,21] also supports global dispersal capacity of GVs. Despite these sporadic studies, no unified analytical framework has yet integrated metagenomic data across global lakes and GV lineages, hindering a comprehensive view of global freshwater GV biogeography.

Compared with marine systems or shallower lakes, deep freshwater lakes are geographically discrete and less susceptible to disturbances [22], making them suitable for examining the role of dispersal limitation in shaping microbial distributions. Moreover, deep lakes exhibit seasonal or persistent thermal stratification, resulting in ecologically distinct water layers [23,24]. This combination of geographic isolation and vertical structure creates a unique framework for investigating biogeography across both vertical and horizontal dimensions. Here, we constructed a genome-resolved view of freshwater GV biogeography across the globe by applying a unified analytical framework to both our original and publicly available metagenomic datasets from deep freshwater lakes spanning five continents. By reconstructing GV metagenome-assembled genomes (MAGs) and profiling their distribution patterns, we resolved their distribution across lakes and water layers at species-level phylogenetic resolution, revealing a complex interplay of factors shaping freshwater GV biogeography.

## Materials and Methods

### Data collection

In this study, we analyzed 203 metagenomic datasets across different water layers from 35 deep freshwater lakes spanning five continents (Australia, Asia, Europe, North America, and Africa) (Fig. S1; Table S1). Among them, 176 are publicly available including 152 short-read samples and 24 long-read samples (Oxford Nanopore) (Table S1). The other 27 samples were collected in this study from six deep freshwater lakes in Japan (Lakes Aoki, Ikeda, Shikaribetsu, Miike, Shikotsu, and Towada) (Table S1). After sample collection, lake water was prefiltered through a 5-µm polycarbonate membrane filter (TMTP14250; Merck Millipore) and then filtered through a 0.22-μm-pore-size Sterivex cartridge (SVGP01050; Merck Millipore) until clogged. The filters were kept in −20 °C until further processing. DNA was extracted from the Sterivex filters (i.e., 0.22- to 5-μm size fraction) using DNeasy PowerSoil Pro Kits (Qiagen) following the manufacturer’s protocol and sequenced by Illumina short-read sequencing platforms (Table S1). In addition, long-read sequencing was performed on four samples from Lake Ikeda (5 m and 100 m) and Shikaribetsu (3 m and 70 m). The long-read sequencing library was prepared using a ligation sequencing kit (SQK-LSK114; Oxford Nanopore) and sequenced by an R10.4.1 flow cell (FLO-MIN114; Oxford Nanopore) using the Oxford Nanopore MinION platform for 72h. Base-calling was performed using Dorado v1.1.1 with sup mode [25].

### MAG reconstruction

For both publicly available and original datasets, Nanopore reads were polished with fmlrc2 v0.1.7 using corresponding short reads and assembled using Flye v2.9.5 (parameters: --nano-corr --meta) [26]. For short-read datasets, quality control and assembly were performed using fastp v1.3.0 [27] (parameters: −3 -W 6 -M 20 -q 15 -u 50 -n 0 -p -l 50) and SPAdes v4.1.0 [28] (parameters: -meta -k 21,33,55,77,99,127), respectively. Contig abundance profiles of all 203 assemblies, including both nanopore- and short-read-derived assemblies, were generated by mapping quality-controlled short reads to assemblies using bwa-mem2 v2.2.1 [29]. For each assembly, reads from samples collected at the same lake were cross-mapped to improve differential coverage signals for binning, with abundance profiles typically derived from two or more samples per assembly. Contig coverage across samples was estimated using jgi_summarize_bam_contig_depths [30], and the resulting multi-sample coverage matrix was provided to MetaBAT2 v2.15.15 [30] for binning.

To identify GV metagenome assembled genomes (MAGs), we first excluded prokaryotic bins based on completeness estimates by CheckM v1.2.5 [31] using bacterial and archaeal marker sets. Bins with estimated completeness >15% (bacterial markers) or >20% (archaeal markers) were classified as prokaryotic and removed from subsequent analyses. For GVs under the phylum *Nucleocytoviricota*, we then performed gene predictions using prodigal-gv [32,33] and used “ncldv_markersearch” tool [2] to screen bins for seven conserved marker genes: *Jelly-roll MCP, PolB, SFIIB, A32, VLTF3, TopII*, and *RNAPL*. Bins containing at least one marker gene were assigned a weighted score based on 20 core genes of nucleocytoviruses, and those with scores >5.75 were retained [34]. For the GV phylum *Mirusviricota*, genes were annotated by prodigal-gv and bins were screened for the presence of mirusvirus *HK97 MCP* gene by hmmsearch using a customized hmm model [5] (bit score > 100). Bins were considered putative mirusvirus MAGs if at least one copy of *HK97 MCP* gene was present.

### Quality control, dereplication, and taxonomic assignments

After decontamination of the contigs (see Suppl. Methods), completeness and contamination of nucleocytovirus MAGs were estimated using gvclass v1.0.0 [35] (see Suppl. Methods). After quality control, dereplication for high- and medium-quality MAGs was performed at 95% average nucleotide identity (ANI) using dRep v3.5.0 (-- ignoreGenomeQuality) [36]. Taxonomic classification of nulceocytoviruses at the order and family levels was performed using TIGTOG [37], and assignments were refined manually based on the phylogenetic topology (see Suppl. Methods). For mirusviruses, taxonomic orders were assigned based on the tree using HK97 MCP sequences of recovered MAGs and references from a recently curated database [4] (see Suppl. Methods).

### Abundance profiling and abundance-based biogeography

We mapped all quality-controlled metagenomic short reads to the collection of high-quality GV MAGs using bwa-mem2 [29] and mapped reads per kilobase of genome per million reads of metagenome (RPKM) and was calculated using CoverM v0.7.0 (genome -- min-read-percent-identity 92 --min-covered-fraction 0.5) [38]. The presence of a MAG in a sample was defined as at least 50% of the genome being covered by the mapped reads. MAGs with a covered fraction below 50% were considered absent in the sample and RPKM values were therefore set to zero in the output. We used this conservative threshold to minimize false-positive detections, while recognizing that it may underestimate the geographic breadth of GV species. Given the strong seasonal dynamics of GV communities in the surface of lakes [5] and the limited sampling time points for each lake, their geographic breadth is inevitably underestimated. Therefore, throughout the study we focused on discussion the wide distributions of GVs rather than their endemism for a reasonable interpretation.

Based on the Bray-Curtis dissimilarities between samples collected during manually identified stratified periods (n=153) in the RPKM profile table, non-metric multidimensional scaling (NMDS) ordinations were generated using the R package vegan [39,40], and environmental parameters manually curated from the associated publications were fitted onto the ordination using the envfit function. Pairwise geographic distances between sampling locations were calculated using the Haversine formula implemented in the geosphere R package. Distance-decay relationships among GV communities were tested by correlating pairwise Bray-Curtis dissimilarity with geographic distance using Spearman’s rank correlation through Mantel tests. For comparison with marine GV communities, metagenomic reads and GV MAGs from a public global marine dataset [17] were reanalyzed using the same analytical pipeline applied to our freshwater dataset.

### Vertical habitat preference assessment

Vertical habitat preference of MAGs was evaluated using samples collected during manually identified stratified periods (n=153). When multiple samples were available within a water layer in the same lake (e.g., due to multiple sampling time points or depths), RPKM values were averaged for each MAG within the water layer. The resulting lake-wise RPKM values were then summed across lakes for each water layer (RPKM_epi_ and RPKM_hypo_). To quantify vertical distribution, we calculated a vertical distribution index (VDI) as:

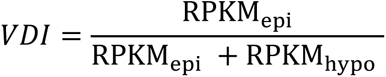

MAGs with an VDI > 0.95 and < 0.05 were classified as epilimnion- and hypolimnion-specific, respectively. Otherwise, the MAGs were considered non-specific, unless they were absent from all stratified samples, where they were designated as unknown.

## Results

### Catalog of GV MAGs across global deep freshwater lakes

Using the metagenomic datasets from 35 lakes across five continents, we recovered 1630 nonredundant nucleocytovirus MAGs after dereplication, including 1126 high-quality MAGs (Table S2). Among these 1630 MAGs, 116 consisted of a single contig, of which 14 harbored terminal inverted repeats (TIRs) with lengths ranging from 127 bp to 34393 bp (Table S3) and 11 were circular genomes (Table S2). The recovered MAGs spanned five orders of the phylum *Nucleocytoviricota*: *Imitervirales* (n=1352), *Pimascovirales* (n=143), *Asfuvirales* (n=60), *Algavirales* (n=43), and *Pandoravirales* (n=29) (Fig. 1A). Genome size of these MAGs ranged from 100.7 to 891.3 kbp and GC content varied from 17.6% to 68.4% (Fig. S2). Using a 95% ANI threshold for species-level clustering, ∼16% (n=261) of the 1630 MAGs clustered with the largest and most up-to-date curated database of GV MAGs currently available [41] (Table S4; Fig. 1C). Those previously described species detected in our dataset were mostly freshwater-derived MAGs (n=255), with several species from other ecosystems such as brackish waters (n=5), saline lakes (n=1), and deep shales (n=2).

**Figure 1.**
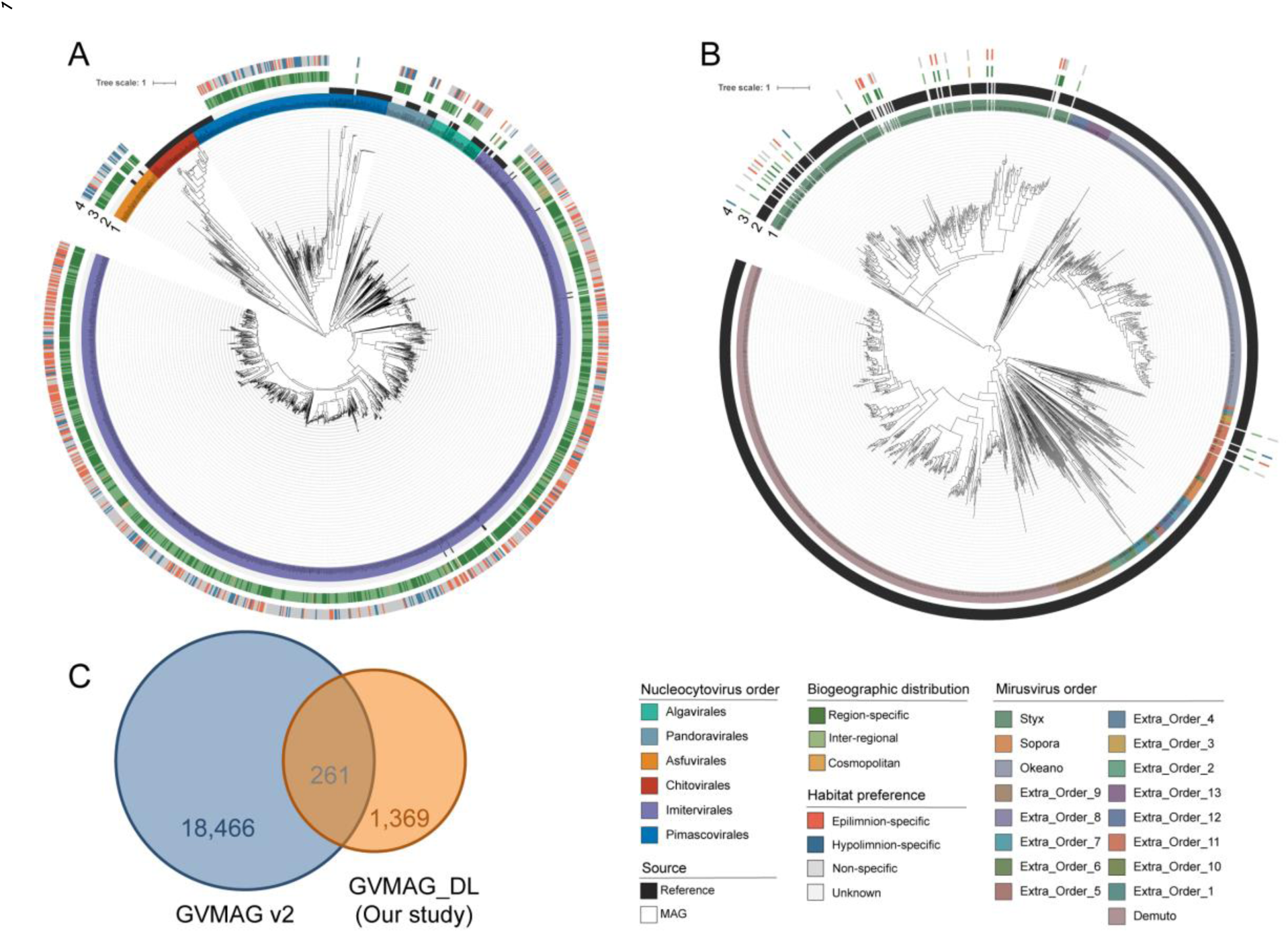
Phylogenetic diversity and novelty of giant virus metageome-assembled genomes (GV MAGs). (A) Phylogenetic tree of the recovered nucleocytovirus MAGs. The tree was built using the concatenated protein sequences of seven genes (*PolB, TFIIB, TopoII, A32, SFII, VLTF3,* and *RNAPL*) with LG+F+I+R10 model and is rooted between the class *Pokkesviricetes* and *Megaviricetes*. (B) Phylogenetic tree of the mirusvirus MAGs using HK97 MCP sequences. Sequences from a recent global survey of *Mirusviricota* MCPs [4] and freshwater mirusvirus MAGs recovered in this study are included. The tree was built using IQ-TREE with the LG + F + R10 model and rooted between Styxvirales and the rest (see Suppl. Methods). For both (A) and (B), the first outer layer of the tree (from the inside) indicates the taxonomic order of each GV genome, including our GV MAGs and the reference GV genomes. In the second layer, the tips colored in black are the reference sequences used to guide taxonomic assignment of our GV MAGs. The third layer indicates the global distribution of our GV MAGs classified by number of detected regions. Region-restricted MAGs were confined to a single region, while inter-regional and cosmopolitan GV MAGs were detected in two to three and four to five regions, respectively. The fourth layer shows the habitat preferences across water layers. GV MAGs are classified into epilimnion- and hypolimnion-specific MAGs or non-specific generalists. Unknown preference means that the habitat preference of this MAG could not be determined using our data (see Methods). Scale bar represents one substitution per site. (C) Overlap between freshwater GV species identified in this study (GVMAG_DL) and the previously established GV MAG database GVMAG V2 [41]. Numbers indicate the count of GV species unique to each dataset and those shared between datasets based on clustering criteria of 95% ANI. The large proportion of unique GV MAGs recovered in this study substantially expands the known diversity of freshwater GVs.

In addition to nucleocytoviruses, we also obtained 33 nonredundant mirusvirus MAGs after dereplication (Table S2). These included 29 MAGs assigned to the order *Styxvirales* and four assigned to a putative order (extra_order_11) proposed in a recent study [4] (Fig. 1B). Through the clustering with the latest mirusvirus genome database [4] at 95% ANI threshold, 12 (36%) MAGs were grouped with previously described MAGs (Table S5). Notably, one mirusvirus MAG (179-MiladaApr19-15_30) was recovered as a single-contig with TIRs, suggesting a complete genome (Table S3).

We adopted the continental names of Europe, Asia, Africa, North America, and Australia to designate regions for GV distributions (Fig. S3). The global distribution of all 1630 nucleocytovirus and 33 mirusvirus MAGs was assessed based on abundance profiles (Fig. S4). Europe and Asia contained the largest count of GV species (n=1467 and 842, respectively), followed by North America (n=299), Africa (n=176), and Australia (n=50). These regional species count likely reflect differences in sampling effort, as Europe and Asia were represented by substantially more lakes and metagenomes than the other regions (Table S1). Europe and Asia shared the largest number of GV species (n=601), accounting for 71.4% of the Asian GV set and 41.0% of the European GV set. The GV species of North America also substantially overlapped with Europe (n=258; 86.3% of North American species) and Asia (n=166; 55.5%). In the following species-level biogeographic analyses, we used only high-quality nucleocytovirus MAGs along with 33 mirusvirus MAGs to ensure robust assessment of the presence/absence and reasonable genomic statistics.

### Geographic prevalence identifies cosmopolitan and region-specific GVs

We evaluated the geographic prevalence of each high-quality nucleocytovirus (n=1126) and mirusvirus MAGs (n=33) based on the number of regions in which it was detected. Nearly half (n=555) of the GVs were identified across multiple regions, while the rest of them (n=604) were region-specific (Fig. 2). The proportion of region-specific MAGs varied across viral orders, with *Pandoravirales* and *Asfuvirales* showing a higher proportion of region-specific members compared to other groups (Fig. 2A).

**Figure 2.**
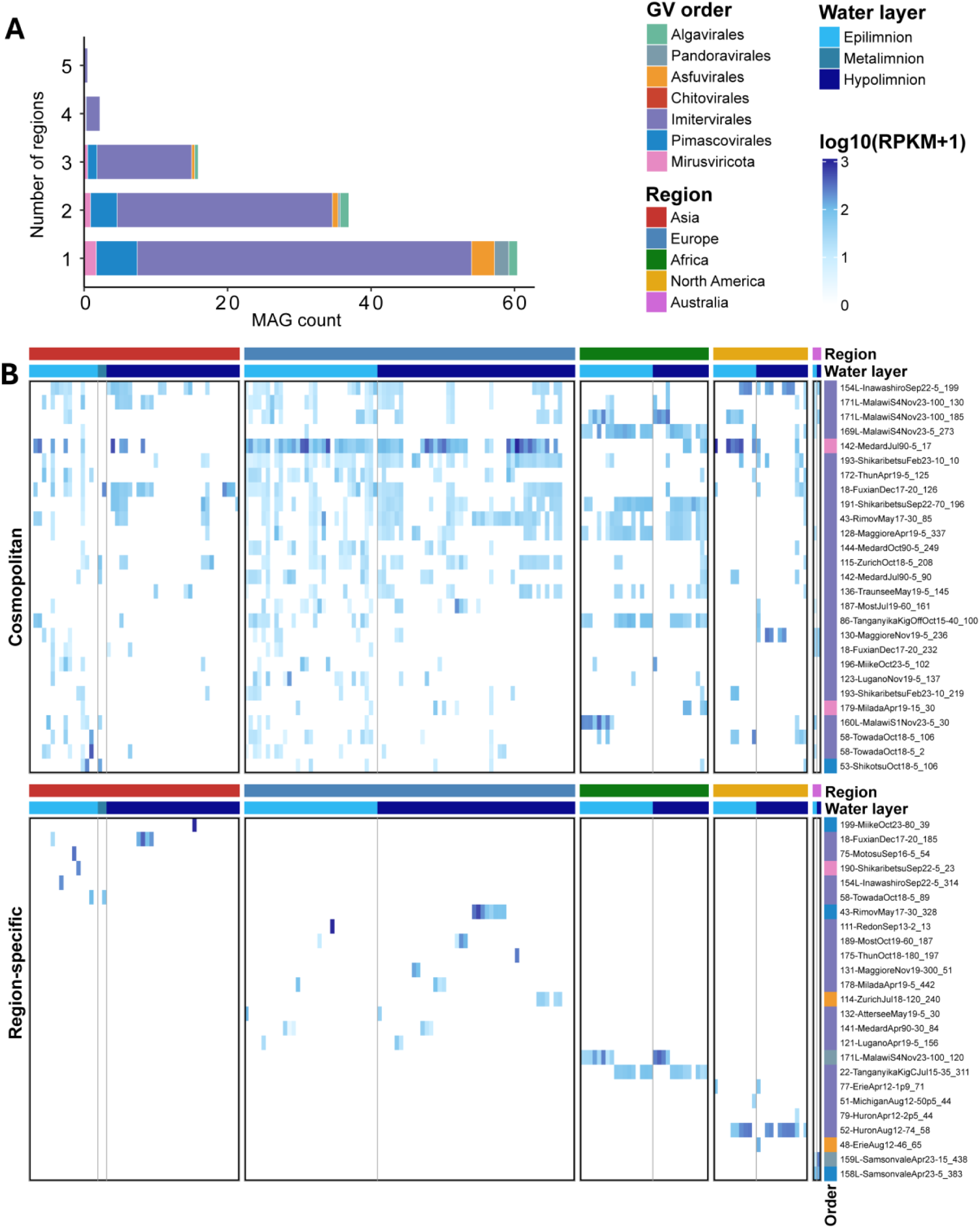
Biogeographic breadth and abundance patterns of freshwater GV MAGs. (A) Distribution of GV MAGs across geographic ranges, where range is defined as the number of regions in which each MAG was detected (see Methods). Bars show MAG counts and are colored by GV order. (B) Heatmap showing the RPKM (reads per kilobases of MAGs per million mapped reads) of 27 cosmopolitan (present in four or five regions) and selected region-specific GV MAGs across freshwater samples. Rows are MAGs and columns are all 203 samples. In total, 27 region-specific MAGs were selected based on abundance rank while maximizing lake representation within each region. Samples are ordered by region first, and then water layer.

Of the 555 GVs distributed across multiple regions, 27 were detected in four or five regions and are hereafter referred to as cosmopolitan GVs (Fig. 2B). The remaining 528 GVs were detected in two or three regions and are designated as inter-regional GVs in this study. Among these inter-regional GVs, 438 (83.0%) were detected exclusively across regions in the Northern Hemisphere, whereas 88 (16.7%) occurred across both hemispheres and only two were shared exclusively among Southern Hemisphere regions. The majority of cosmopolitan GVs (n = 24) belonged to the order *Imitervirales*, whereas one MAG was assigned to *Pimascovirales* (53-ShikotsuOct18-5_106) and two mirusviruses (142-MedardJul90-5_17; complete genome 179-MiladaApr19-15_30) of the order *Styxvirales* (Fig. 2B). Among the cosmopolitan GVs, the *Pimascovirales* MAG and three *Imitervirales* MAGs of the family IM_01 were epilimnion-specific, whereas the remaining MAGs exhibited non-specific habitat preferences and none were hypolimnion-specific. At a finer taxonomic resolution, these cosmopolitan imiterviruses were scattered across several families (Fig. 1A), with 13 members belonging to IM_01, followed by IM_13 (n = 6), IM_16 (n = 3), and IM_12 (n = 2). Notably, all 10 MAGs of family IM_13 recovered in this study exhibited inter-regional or cosmopolitan distributions (Table S2; Fig. 1A). In addition, members of family AG_01 and few distinct clades within family IM_01 were also enriched for MAGs with inter-regional distributions (Table S2; Fig. 1A).

### Vertical partitioning of GVs across freshwater lake ecosystems

The vertical habitat preference of each GV MAG, evaluated by the VDI scores (see Methods), showed bimodal distribution (Fig. 3A), indicating that freshwater GVs exhibit clear vertical habitat specialization. Of the 1120 GV MAGs detected in stratified samples, 312 (27.8%) and 177 (15.8%) were classified as epilimnion- and hypolimnion-specific, respectively. Their vertical habitat preferences were highly consistent, with most MAGs repeatedly detected in the same water layer across different lakes rather than exhibiting lake-specific shifts in depth distribution (Fig. 3B). Notably, one of the most abundant hypolimnion-specific GVs in the dataset (157L-ToyaSep22-150_4 of the family IM_01), was distributed across three regions and 23 lakes. This MAG was consistently enriched in the hypolimnion of multiple Japanese lakes (Lakes Motosu, Shikotsu, Towada, Biwa, Ikeda, Miike, Chuzenji, and Toya), while also occurring at lower abundance in several European and North American lakes (Table S6). In contrast, 53-ShikotsuOct18-5_15 was exclusively detected in the epilimnion of Lakes Motosu and Shikotsu with high abundance. Interestingly, several cosmopolitan GV MAGs (e.g., 160L-MalawiS1Nov23-5_30 and 58-TowadaOct18-5_2) were also exclusively detected in epilimnion despite spanning four and five regions, respectively.

**Figure 3.**
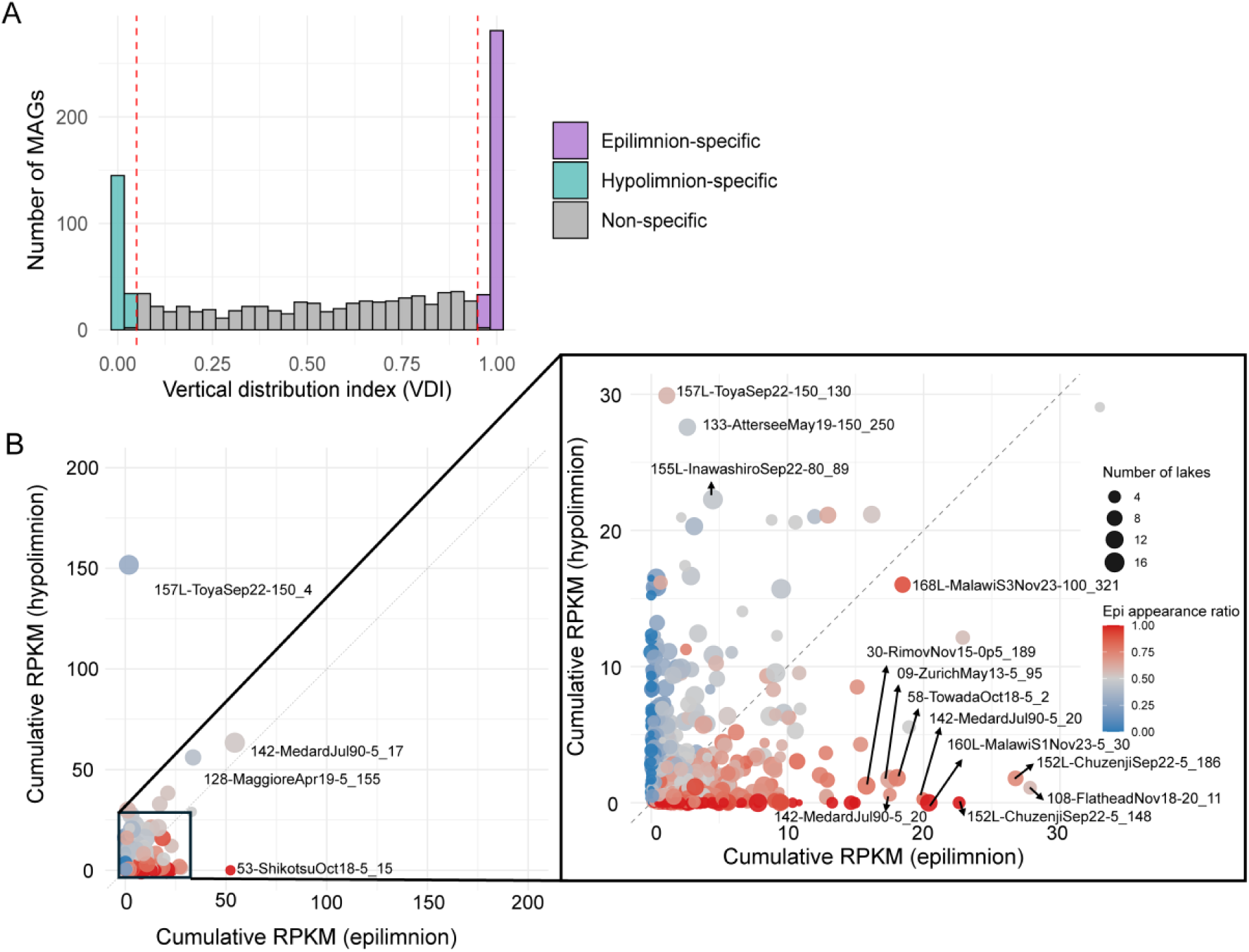
Vertical partitioning of GV species in stratified samples. (A) Histogram of vertical distribution index (VDI). VDI is calculated for each MAG as accumulative lake-level RPKMs in epilimnion samples divided by total lake-level RPKMs across all lakes (see Methods). The threshold of 0.95 and 0.05 was used to determine if a MAG is epilimnion- or hypolimnion-specific. (B) Consistency of vertical partitioning across lakes. Each point represents a GV MAG and its size indicates the number of lakes in which the MAG was detected. Point colors represent the proportion of epilimnion detections among all detections across lakes, with red and blue indicating biased detections in epilimnion- and hypolimnion samples, respectively.

### Community divergences across water layer and regions

Beta-diversity analysis revealed that GV communities from African lakes clustered distinctly from the samples from other lakes (PERMANOVA, *R²* = 0.107, *p* = 0.001) (Fig. 4A). Within Africa, samples from the two lakes (Lake Malawi and Lake Tanganyika) formed individual clusters. Consistently, Bray-Curtis dissimilarity of Africa-involved inter-regional sample pairs appeared to be the highest (Fig. S5A). These patterns likely reflect the dominance of region-specific taxa in African lakes (Fig. S6A). In addition, vertical partitioning of the community structure was less obvious in African lakes (Fig. S5B), consistent with the lowest relative abundances of epilimnion- and hypolimnion-specific GVs (Fig. S6B).

**Figure 4.**
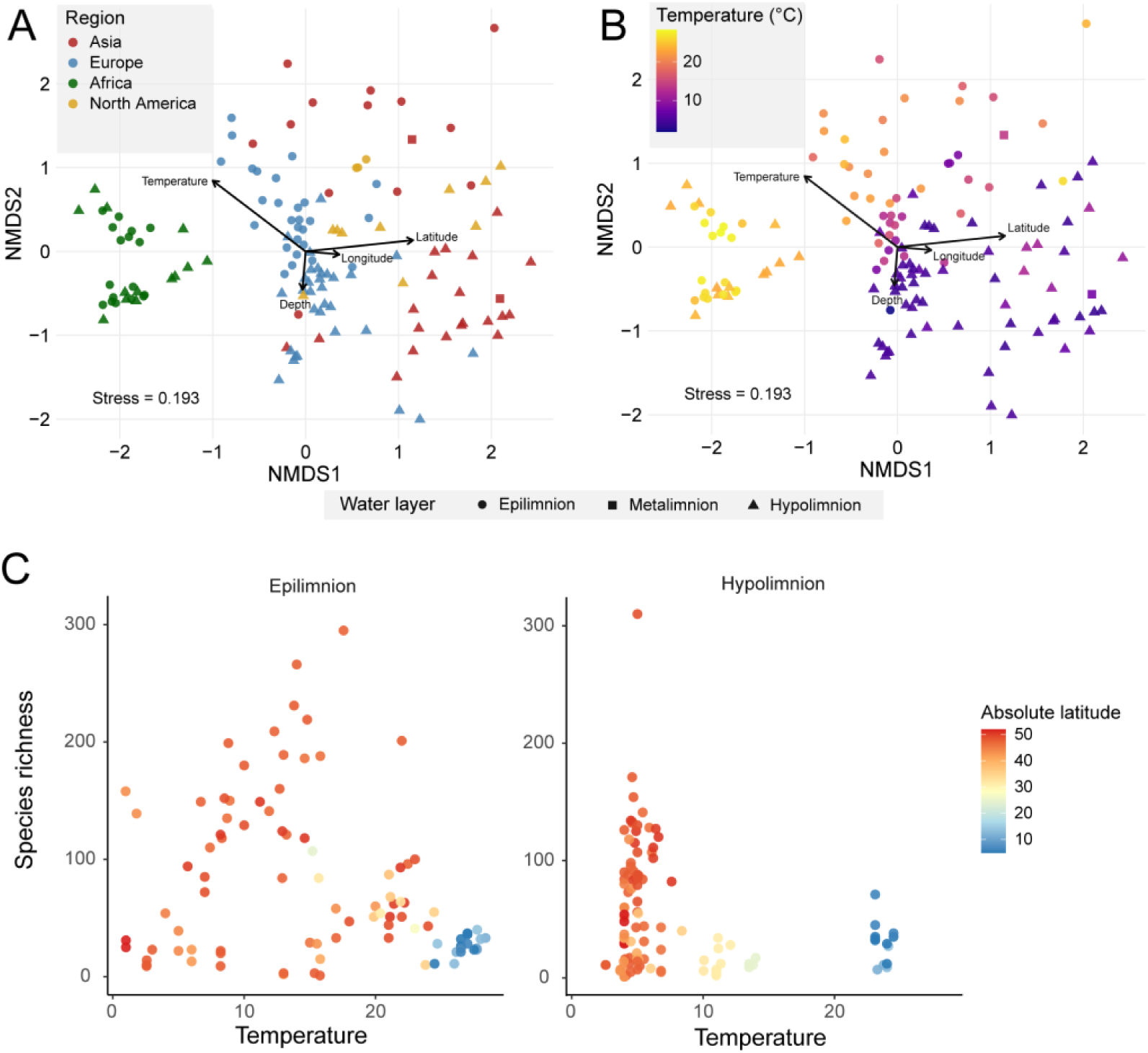
Biogeographic patterns and environmental associations of GV communities in stratified freshwater lakes. (A–B) Non-metric multidimensional scaling (NMDS) ordination of GV community composition across samples from stratified period based on Bray–Curtis dissimilarity. Environmental parameters were fitted onto the NMDS ordination, with significance assessed by permutation testing (n = 999). Temperature (*R²* = 0.64, *p* = 0.001), latitude (*R²* = 0.51, *p* = 0.001), depth (*R²* = 0.08, *p* = 0.007), and longitude (*R²* = 0.05, *p* = 0.033) were significantly associated with variation in community composition. Each point represents a sample, with shapes indicating water layer (epi-, meta-, and hypolimnion). Points are colored by region (A) and water temperature (B). (C) Relationship between GV species richness and water temperature across samples, separated by water layer. Points are colored by absolute latitude of the sample.

In contrast, among the lakes in Europe, Asia, and North America, the epilimnion and hypolimnion samples clustered separately regardless of geographic distance (PERMANOVA, *R²* = 0.046, *p* = 0.001), indicating that the communities are structured not only by region but also by water layers (Fig. 4A, B). Asia showed the highest relative abundance of epilimnion or hypolimnion-specific GVs, followed by Europe (Fig. S6B). Correspondingly, vertical partitioning patterns were the most obvious in the NMDS ordination for these regions, in contrast to the weak partitioning observed in African lakes (Fig. 4B). The strong vertical partitioning of GV communities reflects the distinct temperature regimes in the epilimnion and hypolimnion (Fig. 4B). Furthermore, the association between species richness and water temperature differed markedly between water layers (Fig. 4C). The species richness increased with temperature up to about 10 °C before declining at higher temperatures in the epilimnion, whereas it showed a continuous decrease with increasing temperature in the hypolimnion.

### Distance-decay relationships in freshwater GV communities

Aligned with the regional grouping patterns, GV communities exhibited significant distance–decay relationships, with community dissimilarity increasing with geographic distance across both intra-regional and inter-regional spatial scales (Mantel *r* = 0.139–0.457, *p* < 0.02; Fig. 5). At the intra-regional scale, beta diversity increased with geographic distance in both the epilimnion and hypolimnion, with a stronger relationship observed in the hypolimnion (Mantel *r* = 0.457) than in the epilimnion (Mantel *r* = 0.195). In contrast, inter-regional comparisons showed substantially weaker spatial relationships, particularly in the epilimnion (Mantel *r* = 0.139), where Bray-Curtis dissimilarity remained consistently high across large geographic distances at inter-regional scales. Although hypolimnion communities retained relatively stronger geographic structuring at the inter-regional scales (Mantel *r* = 0.181), community dissimilarity generally converged toward consistently high values with reduced variance compared to intra-regional comparisons, indicating geographic distance explained progressively less variation in community composition at broader spatial scales (Fig. S7). Moreover, a comparison with marine GV communities further revealed markedly stronger distance-decay relationships in freshwater epilimnion communities than in marine surface communities across the full geographic range (freshwater epilimnion: Mantel *r* = 0.502; marine surface: Mantel *r* = 0.122; *p* < 0.001; Fig. S8).

**Figure 5.**
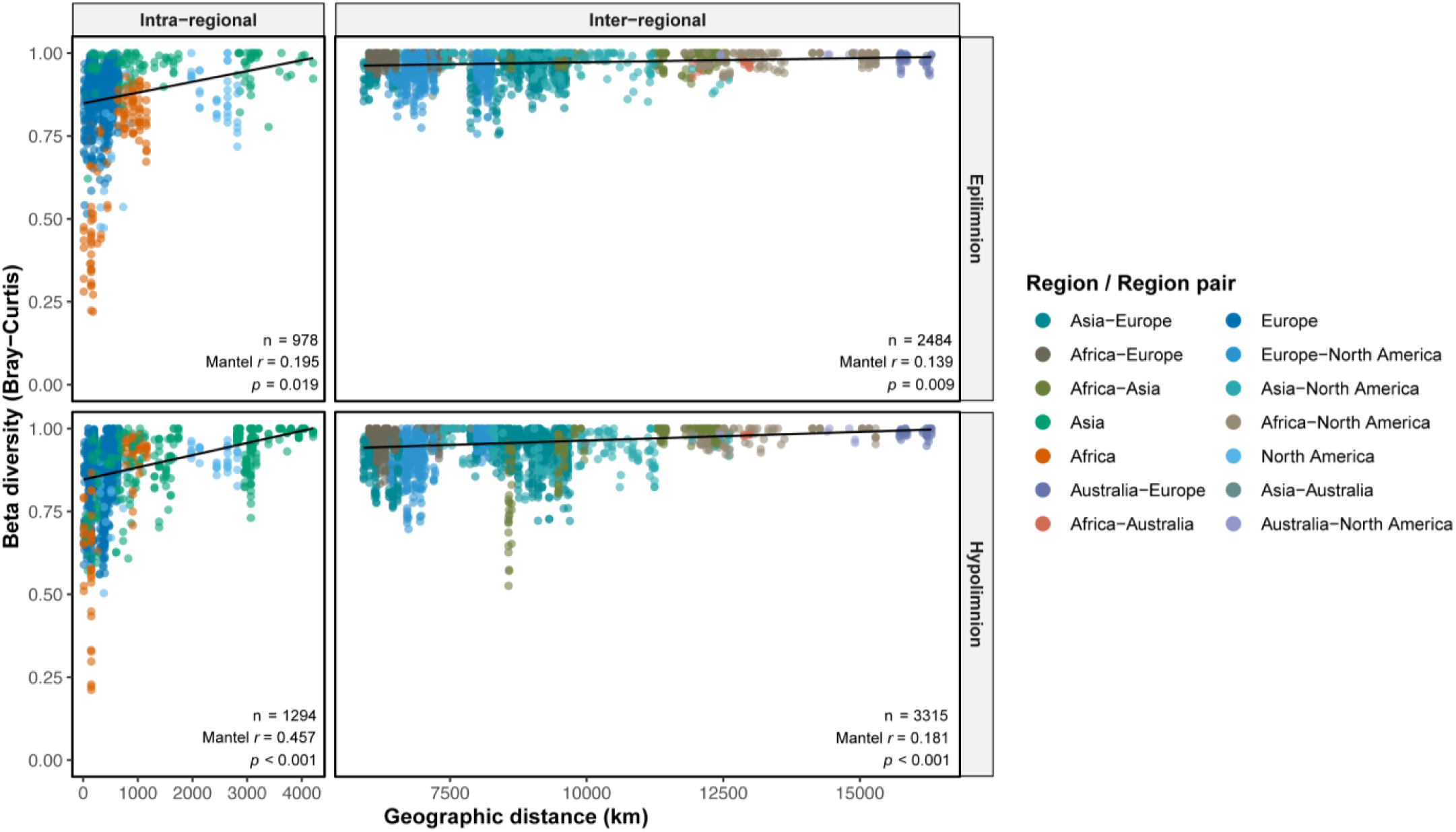
Distance–decay relationships of freshwater GV communities across intra-regional and inter-regional spatial scales. Beta diversity was quantified using Bray-Curtis dissimilarity and plotted against geographic distance for epilimnion and hypolimnion communities. Spearman’s rank correlations assessed through Mantel tests are given on the lower right of each plot. Linear regressions are shown as black lines. These regressions were not used to infer correlations between community dissimilarity and geographic distance but such correlations are quantified by the Mantel test results.

### Broadly distributed GVs possess larger genomes and broader functional repertoires

We observed a positive correlation between GV genome size and their geographic breadth, measured as the number of regions (Spearman’s *ρ* = 0.161, *p* < 0.001; Fig. 6A) and lakes (Spearman’s *ρ* = 0.162, *p* < 0.001; Fig. 6B) in which each MAG was detected. The positive trend was mainly driven by *Imitervirales* as the dominant order in our dataset, whereas other groups showed weak or no correlations (Fig. S9). Consistent with this pattern, the absolute number of auxiliary metabolic genes (AMGs) increased with regional occupancy (Spearman’s *ρ* = 0.133, *p* < 0.001; Fig. 6C), while AMG density normalized by genome size did not (Fig. S10A). Similarly, the number of unique AMG increased with regional occupancy (Spearman’s *ρ* = 0.125, *p* = 2.07 × 10⁻⁵; Fig. 6D), whereas the number of unique AMGs per Mbp remained unchanged (Fig. S10B). We identified 25 orthogroups detected exclusively in cosmopolitan/inter-regional MAGs, the majority of which lacked functional annotation (Table S7). Among the annotated OGs were saposin-like proteins (K12382) and CS-domain proteins.

**Figure 6.**
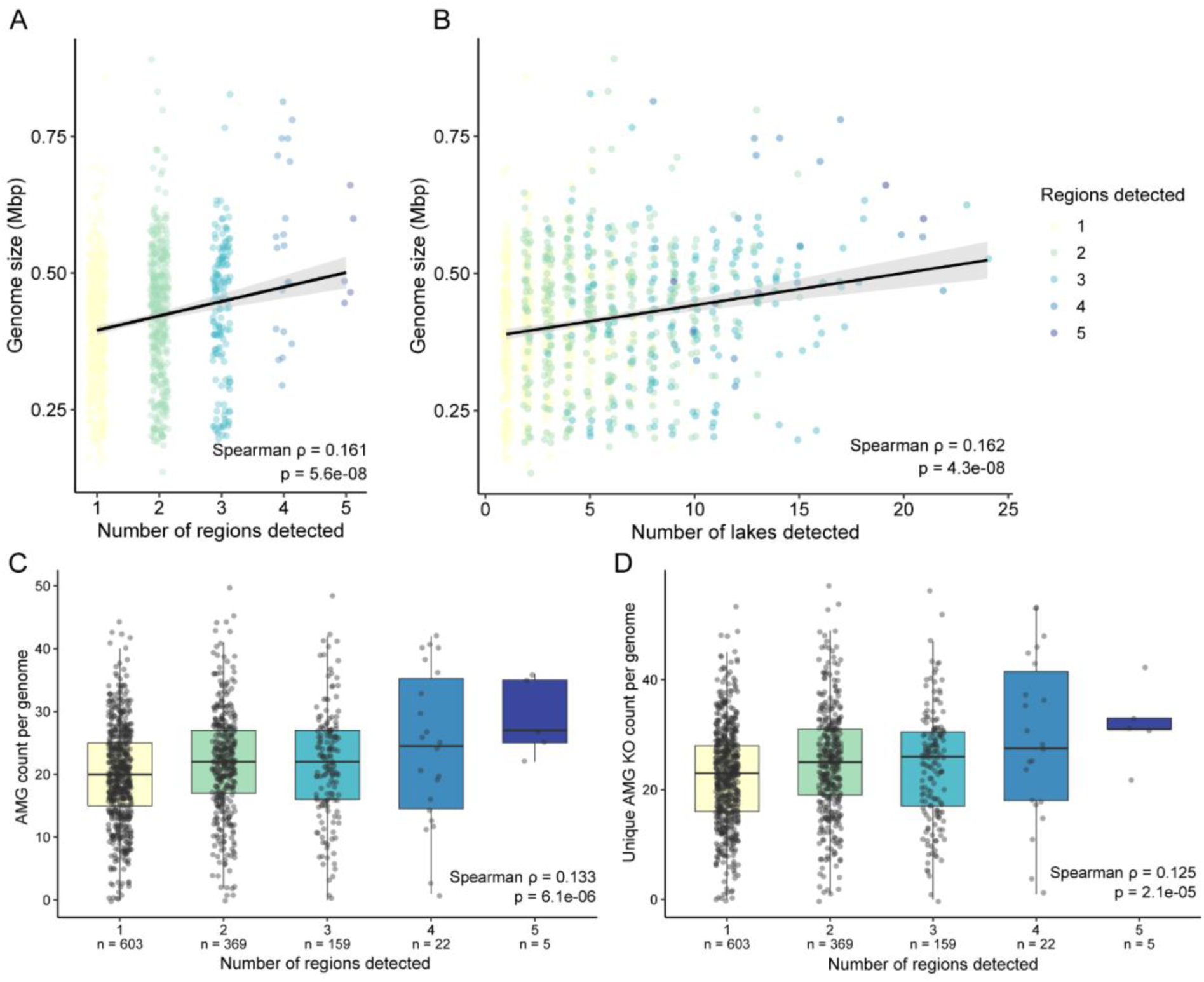
Associations between geographic occupancy, genome size, and AMG repertoire. (A–B) Relationships between GV genome size and geographic occupancy, measured as the number of detected regions (A) and lakes (B). Each point represents a high-quality GV MAG and is colored according to the number of detected regions. Black lines indicate linear regression fits with 95% confidence intervals shown in gray. Spearman’s rank correlation indicates a significantly positive correlation both in (A) and (B) with coefficients (ρ) of 0.161 and 0.162, respectively. (C–D) Distributions of total AMG abundance (C) and unique AMG KO counts (D) across GV MAGs grouped by the number of regions in which they were detected. Individual points represent MAGs, and sample sizes for each occupancy category are shown below the x-axis. Spearman’s rank correlation indicates a significantly positive correlation both in (C) and (D) with coefficients (ρ) of 0.125 and 0.133, respectively. Genome size, AMG abundance, and AMG functional diversity all increased significantly with geographic occupancy.

## Discussion

### A global genome catalog reveals the coexistence of cosmopolitan and endemic GV lineages

By integrating both long- and short-read metagenomic datasets from 35 deep lakes spanning five continents, including underexplored regions in the Southern Hemisphere and six newly sampled Japanese lakes, we established the geographically most comprehensive catalog of GV genomes in deep lakes to date and substantially expanded the known diversity of freshwater GVs (Fig. 4C). Despite the growing number of freshwater GV reference databases (n=4957) [41], only 16% of the GV MAGs recovered in this study clustered with previously described species, most of which were originally identified from freshwater environments (Table S4; S5). Our dataset systematically incorporates both epilimnion and hypolimnion GV communities from deep lakes. The extensive novelty of our GV MAGs suggests that deep lake ecosystems harbor a large reservoir of GV diversity that remains poorly represented in the current genome catalogs.

Leveraging this global genome catalog, we resolved the GV diversity at species-level resolution and revealed that biogeographic breadth is highly heterogeneous among GV lineages. All GV orders contained both geographically restricted and broadly distributed lineages (Fig. 2A). Previous biogeographic investigations primarily based on hallmark genes often emphasized the global distribution of GVs [1,13], whereas genome-resolved analyses increasingly reveal endemic GV MAGs in marine and freshwater ecosystems [14,15,19]. Our species-level investigation supports the coexistence of cosmopolitan and endemic GV lineages in local communities. At one extreme, we identified 203 GV MAGs that were detected exclusively within a single lake (Table S2), suggesting high endemism. At the other extreme, five MAGs were identified as cosmopolitan, being detected in all five regions spanning both North and South Hemispheres. These contrasting patterns therefore argue against a strict interpretation of the Baas Becking hypothesis (“Everything is everywhere”) [42].

### Horizontal biogeographic structuring reflects ecosystem connectivity

Despite occasional long-distance dispersal of individual GV lineages, freshwater GV communities exhibited strong regional structuring. Distance-decay analyses showed that community dissimilarity increased strongly at intra-regional scales, whereas the beta-diversity remained consistently high (median=0.98) at inter-regional distances (Fig. 5; Fig. S7), indicating substantial compositional differentiations among geographically distant lake ecosystems. This pattern has also been observed in marine microbial communities, where community dissimilarity remains consistently high across ocean basins, such that geographic distance explains little additional variation and environmental gradients become the primary determinants of community composition [10,43]. Our comparison using identical analytical framework showed that freshwater surface GV communities exhibited steeper distance-decay relationships than marine surface communities (Fig. S8). This finding suggests a stronger dispersal limitation among freshwater ecosystems, likely reflecting the spatially discrete nature of lakes in contrast to the physically interconnected marine environments, which may restrict long-distance viral dispersal and promote local diversification within individual lakes. Of note, pairwise ANI comparisons among MAGs prior to dereplication revealed only six inter-regional genome pairs exceeding 97% ANI with >80% aligned fractions (Fig. S11), suggesting that although some GV lineages are capable of inter-regional dispersal, such events remain rare even among cosmopolitan species. Additional support for the role of ecosystem connectivity comes from comparisons between water layers. Hypolimnion GV communities exhibited significantly stronger distance–decay relationships than epilimnion communities (Fig. 5), consistent with the greater ecological isolation and environmental stability of the hypolimnion [22]. Similarly, global ocean surveys have shown stronger horizontal biogeographic differentiation of microbial plankton communities at deep water layers than surface [9]. Together, our results support the idea that the strength of microbial distance-decay relationships scale inversely with ecosystem connectivity across aquatic environments.

### Vertical partitioning of GV communities and environmental selection

We found that nearly half of recovered GV MAGs showed habitat preference for either epilimnion or hypolimnion (Fig. 3A). By analyzing the global lake samples, we show that vertical water-layer preferences are largely conserved across geographically distant lakes (Fig. 3B). Moreover, the habitat preferences of each GV inferred from our global dataset were highly concordant with those previously reported for GV MAGs collected through spatiotemporal sampling (bilayer for 12 months) in Lake Biwa [5]. Among the 28 GV species identified as water layer-specific in both Lake Biwa and this study, 27 showed consistent water layer preferences (Table S8). This rather consistent habitat preference across lakes and seasons suggest that vertical habitat preference is an intrinsic property of many freshwater GV lineages. Furthermore, the bimodal distribution of vertical habitat preferences remained consistent regardless of geographic breadth (Fig. S12), suggesting that vertical zonation is a conserved ecological property. Aligned with this, both water layer-specific and non-specific MAGs were observed among broadly distributed lineages (Fig. 3B), supporting the idea that geographic distribution and vertical breadth are independent. In deep freshwater lake ecosystems, similarly widespread but hypolimnion-specific taxa have been documented for both bacteria [44,45] and microbial eukaryotes [46,47]. Beyond freshwater lakes, vertical partitioning has also been observed for marine GV communities inhabiting distinct ocean water layers [16,17]. Consistent with these observations, our results suggest that broad geographic distributions do not necessarily imply ecological generalism. Freshwater GV biogeography reflects the interplay between long-distance dispersal and vertical zonation.

Consistent with species-level vertical partitioning, NMDS ordinations repeatedly revealed clustering of GV communities between epilimnion and hypolimnion across numerous non-African lakes (Fig. 4). Such vertical partitioning of GV communities develops with the thermal stratification in deep lakes, which is accompanied by multiple co-varying physicochemical gradients. Among the environmental variables, water temperature appeared to be one of the important factors associated with the vertical clustering of GV communities (Fig. 4B). We therefore further examined temperature-dependent patterns within each water layer and found contrasting relationships between GV richness and temperature (Fig. 4C). These contrasting richness patterns indicate that thermal conditions influence vertically partitioned GV communities in different ways. The richness peak in epilimnion communities coincided with water temperatures representing the spring and autumn blooms commonly observed in temperate lakes (Fig. 4C; Table S9), suggesting that temperature may indirectly shape the GV diversity by driving seasonal host blooms. In contrast, hypolimnion communities showed progressively lower richness in warmer lakes, suggesting the importance of cold, thermally stable deep-water environments for maintaining GV diversity in the hypolimnion.

Freshwater GV richness increased toward higher latitudes in both epilimnion and hypolimnion communities (Fig. S13), contrasting with the classical latitudinal diversity gradient of tropical diversity maximum observed for many macroorganisms [48] and planktonic microeukaryotes [49,50]. This observation suggests that the factors shaping freshwater GV diversity extend beyond host diversity. A similar inverse latitudinal pattern has recently been reported for marine GVs [15,51], where possible explanations for this apparent decoupling from host diversity have been proposed as temporal seasonality of sampling effects and the potentially broader host ranges of GVs enabled by phagocytosis-mediated host entry. Likewise, marine RNA viruses also exhibit diversity patterns largely decoupled from those of their eukaryotic hosts, a pattern that has been suggested to reflect extensive host sharing and highly connected virus–host interaction networks [52]. Our analyses also indicate that thermal regimes likely play an important role in shaping GV richness in the hypolimnion layers (Fig. 4), likely through their effects on closely associated hosts inhabiting the cold, thermally stable environments of deep lakes. It is noteworthy that these deep-water habitats are particularly vulnerable to climate-driven warming, resulting in thermal structural changes of the water column [53]. The GV communities in the hypolimnion may become increasingly vulnerable to biodiversity loss as climate warming progressively alters deep-lake thermal structure.

### Evolution emerges as a third dimension of freshwater GV biogeography

Our results further suggest that freshwater GV biogeography is shaped by lineage-specific evolutionary characteristics. Specifically, several GV lineages, including subclades in the family IM_01, IM_13, AG_01, and PM_01, were enriched among widely distributed GVs (Table S2). This lineage-dependent pattern suggests that intrinsic biological characteristics contribute to differences in biogeographic breadth. Among these lineages, the broad distribution of IM_01 viruses may reflect their relatively wide host range, as members of this family have been linked to diverse and highly abundant eukaryotic hosts across aquatic ecosystems (e.g., haptophytes in marine environments and chrysophytes in freshwater lakes) [54,55]. In contrast, the hosts of the other cosmopolitan-enriched lineages remain poorly resolved. PM_01 viruses have been proposed to associate with haptophyte algae based on co-occurrence analyses [19], although their natural hosts remain largely unknown. Likewise, the host associations of freshwater AG_01 and IM_13 viruses remain unexplored. Identification of their hosts will be a critical subject of future work for understanding the mechanisms underlying their broad geographic distributions.

In addition to the lineage specificity of GV biogeography, we found that widely distributed GVs tended to possess larger genomes and broader functional repertoires than geographically restricted lineages using only high-quality MAGs (Fig. 6). Broadly distributed GVs were also enriched in genes associated with host interaction and intracellular manipulation (Table S7), including saposin-like proteins and CS-domain proteins. Saposin-like proteins are membrane-active lipid-binding proteins involved in membrane-disruptive and host cell interaction processes through lipid-binding activities in microbial eukaryotes [56–58]. The enrichment of these proteins in GV genomes (Table S7) with wide distributions suggests that similar membrane interaction capabilities may enhance viral host entry and infection success across diverse environments. CS-domain proteins contain conserved protein-interaction modules that frequently mediate HSP90 co-chaperone activity in eukaryotic protein-folding and regulatory complexes [59,60]. The enrichment of these functions in widely distributed GVs suggests that enhanced capacities for host interaction and intracellular manipulation may contribute to ecological versatility across diverse environments. Consistent with previous studies, larger GV genomes often encode expanded repertoires of metabolic genes, host-interaction functions, and transcriptional machinery [1,54,61], potentially increasing ecological flexibility through enhanced host attachment capacity and broader host range [54,62–64]. Collectively, our data support that genome expansion may facilitate the persistence and dispersal of certain GV lineages across geographically and environmentally distinct freshwater ecosystems. Such genomic traits are recognized as an important factor determining the biogeography of GVs in deep lake ecosystems.

### Unique biogeographic patterns in African lakes

We found that African lakes exhibited distinct biogeographic patterns characterized by stronger geographic isolation but weaker vertical partitioning than lakes from other regions (Fig. 4A, B; Fig. S5). This unique pattern may reflect the exceptional limnological history of the African Great Lakes, which are among the oldest and most geographically isolated freshwater ecosystems on Earth [65,66]. Similar biogeographic patterns have recently been reported for freshwater SAR11 bacterioplankton [67,68], where ancient lake history may contribute to lineage diversification and population structure beyond dispersal alone. The weak vertical partitioning of the GV community may also be attributed to passive downward transport of virions from surface waters, as previously proposed for marine GVs [13,16]. This process may be particularly important in African lakes with permanently anoxic hypolimnion layers [69]. In particular, Lake Tanganyika exhibited even weaker across-layer community differentiation than Lake Malawi (Fig. S5B), consistent with the contrast in oxygen conditions between its anoxic hypolimnion below 100 m depth and the largely oxygenated sampled water column of Lake Malawi (5–200 m). Anoxic environments often support reduced or compositionally distinct eukaryotic communities relative to oxygenated waters [70,71], while many known or predicted GV hosts, including photosynthetic algae and aerobic protists, are strongly constrained by oxygen [13,55,72,73]. Together, these processes may explain why a larger fraction of GVs in African lakes exhibit no specific preference for either water layer (Fig. S6A). Notably, tropical and deep anoxic lakes remain underrepresented in our datasets. Broader depth-resolved sampling across tropical lake ecosystems will therefore be essential to determine whether the weak vertical partitioning is general feature among tropical freshwater ecosystems.

### Conclusions

We revealed that freshwater GV biogeography is structured along three interacting dimensions: horizontal geographic dispersal and colonization, conserved vertical partitioning, and lineage-specific evolutionary characteristics. The coexistence of cosmopolitan and endemic GV lineages demonstrates that dispersal and colonization capacities vary markedly among lineages, whereas the conservation of vertical habitat preferences across globally distributed lakes underscores its fundamental importance in freshwater GVs. Moreover, the enrichment of several GV lineages (e.g., subclades within the families IM_01, IM_13, AG_01, and PM_01) among broadly distributed GVs, together with the association of broader geographic distributions with genome expansion, suggests that evolutionary innovation may contribute to ecological versatility and long-distance persistence across freshwater environments. Our findings establish a multidimensional framework for understanding freshwater GV biogeography but also highlight that the host ecology of many cosmopolitan GV lineages remains largely unknown. Resolving the interactions between these viruses and their hosts will be key to understanding the mechanisms behind their global distributions.

## Data availability

The nucleotide and protein sequences of the nonredundant GV MAGs, along with alignment and tree files for the phylogenetic analyses conducted in this study, are available at GenomeNet: https://www.genome.jp/ftp/db/community/DeepLakesGVMAGs/. The raw Illumina and Oxford Nanopore reads generated in this study are available under BioProject ID PRJDB42766.

## Acknowledgements

This work was supported by the Japan Society for the Promotion of Science (JSPS KAKENHI Grant Numbers JP18J00300, JP19H03302, JP22H00384 and JP22H00382), the Japan Science and Technology Agency (JST FOREST; Grant Numbers JPMJFR2273 and JPMJFR231C), the Kyoto University Foundation, the Kyoto University Division of Graduate Studies (DoGS) SPRING Program and the Collaborative Research Program of the Institute for Chemical Research, Kyoto University (2024-28, 2025-27). Computational resources were provided by the Supercomputer System of the Institute for Chemical Research, Kyoto University. This study also benefited from the Joint Research Program of the Center for Ecological Research, Kyoto University and Advanced Genomics Center at the National Institute of Genetics. MMS was supported by the Czech Science Foundation (grant 22-03662S). We thank Dr. Shang Shen for assistance with field sampling at Lake Miike and the Tokachi Shikaoi Geopark Promotion Council for supporting field sampling at Lake Shikaribetsu. We are also grateful to Dr. Adrian-Stefan Andrei for valuable comments on the manuscript, and Dr. Russell Young Neches for insightful discussions.

