## Supplementary file for "Genome-resolved biogeography reveals multidimensional structuring of freshwater giant viruses across global deep lakes"

### This PDF file includes:

Figures S1 to S13

Captions for Table S1 to S9

Supplementary Methods

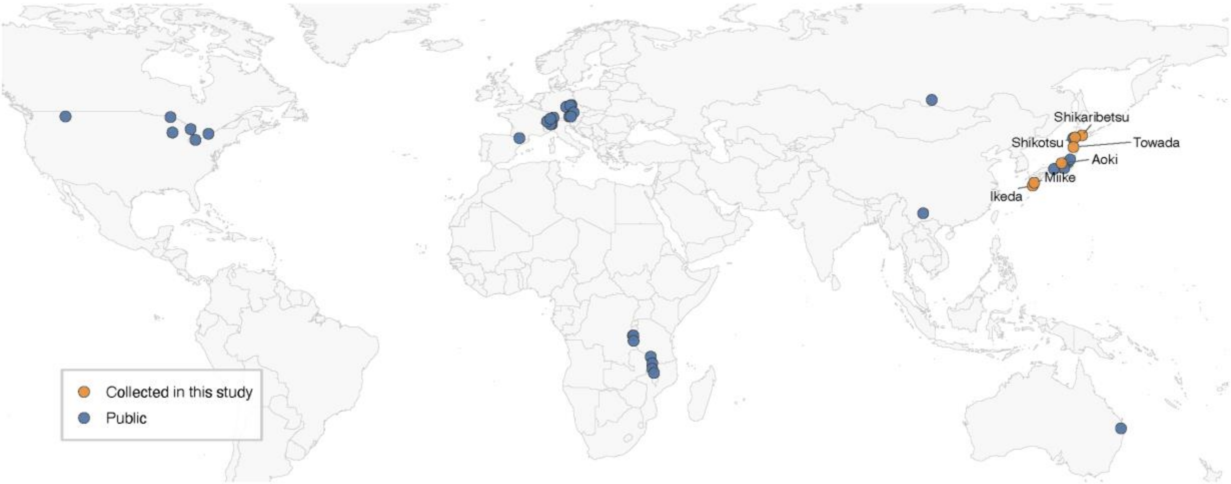


**Supplementary Figure 1. Global map of sampling sites.** A total of 203 metagenomes from 35 deep freshwater lakes spanning five continents were included in this study, comprising 176 publicly available metagenomes and 27 metagenomes collected from six Japanese lakes in this study. Detailed sample information is provided in Table S1.

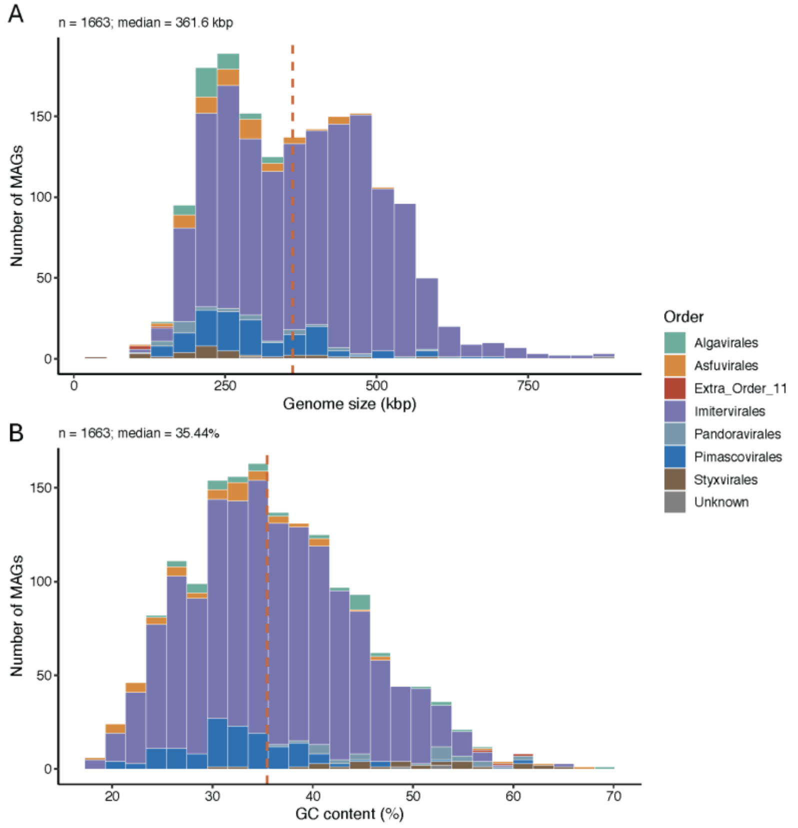


**Supplementary Figure 2. Histogram of genome sizes (A) and GC content (B) of GV MAGs.** The histogram bars in both (A) and (B) are colored by the GV orders. The dashed line indicates the median of genome size or GC content.

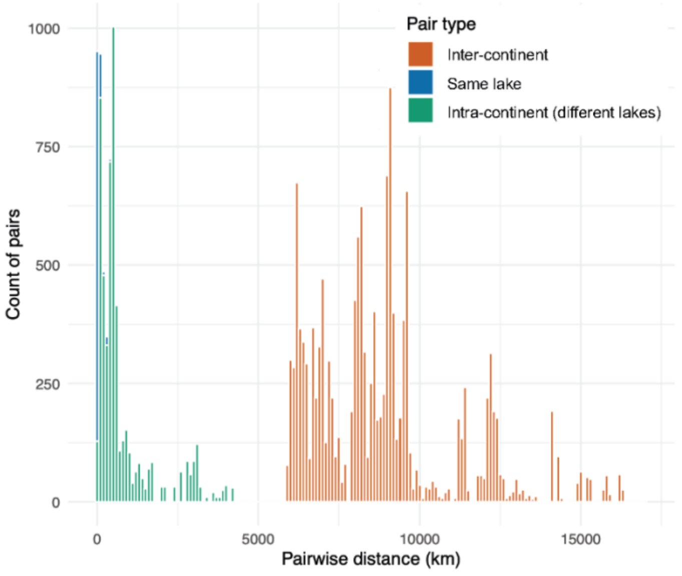


**Supplementary Figure 3. Pairwise distance distribution of all samples.** We identified a natural gap at approximately 5,000 km in the distribution of pairwise geographic distances among samples, which effectively corresponded to continental boundaries. We therefore designated the resulting regions by continent (Europe, Asia, Africa, North America, and Australia) and used them to distinguish intra-regional and inter-regional GV diversity patterns.

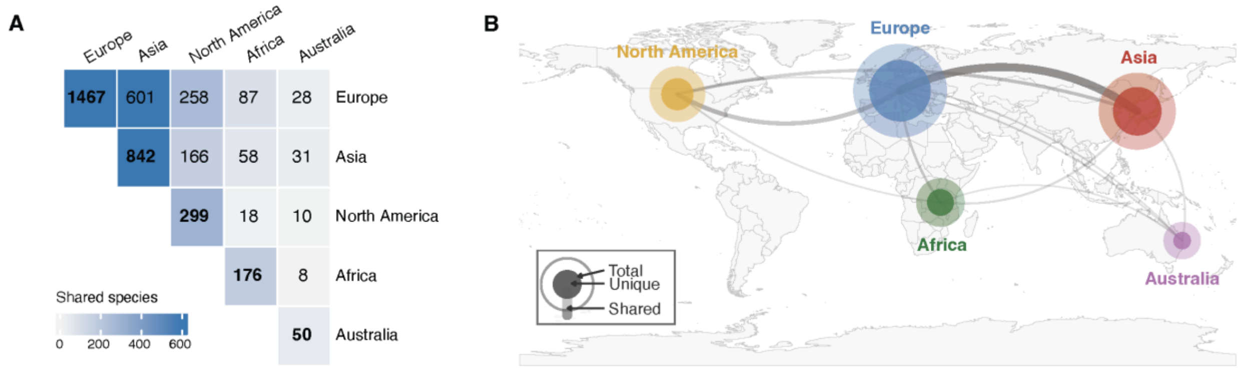


**Supplementary Figure 4. Global distribution of GV species.** (A) Upper-triangular heatmap showing the number of shared GV species between each pair of regions. Diagonal cells indicate the total number of nonredundant GV species detected in each region. (B) Nonredundant region-level GV species richness. Outer circles indicate the total number of nonredundant species detected in each region, whereas the inner circles represent region-specific species. Curved links represent pairwise shared species between regions, with line width and opacity scaled to the number of shared species.

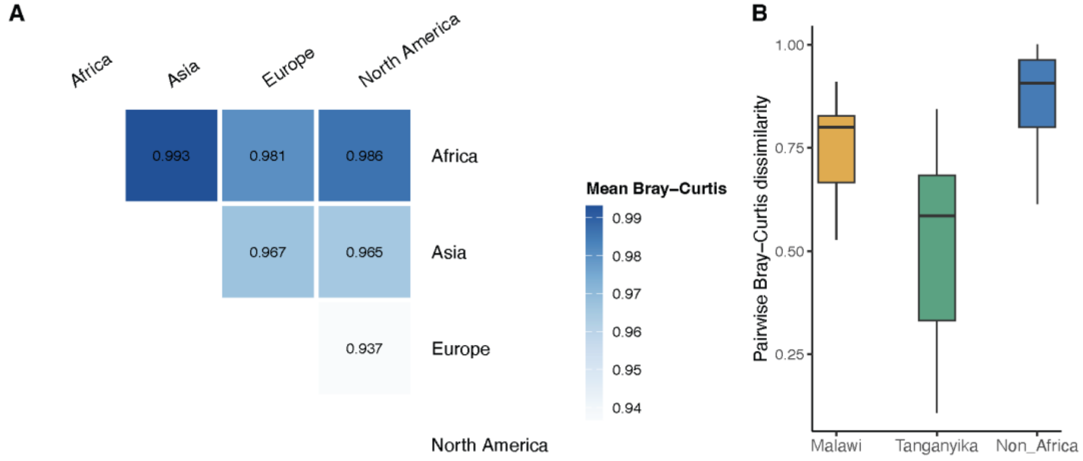


**Supplementary Figure 5. Comparison of biogeographic and vertical patterns in GV beta diversity across regions.** (A) Average Bray-Curtis dissimilarity among across-region sample pairs. The analysis included only samples with ≥5 species that were collected during the stratified period. (B) Bray-Curtis dissimilarity between epi- and hypolimnion GV communities within the same lake, comparing African (Lake Malawi and Lake Tanganyika) and non-African lakes (Australian lakes excluded). Analyses were restricted to stratified samples assigned to either epi- or hypolimnion, and samples with less than five detected MAGs were excluded. Non-African lakes showed the highest dissimilarity (n = 147 pairs; median = 0.907), followed by

Lake Malawi (n = 32 pairs; median = 0.800), whereas Lake Tanganyika showed the lowest dissimilarity (n = 80 pairs; median = 0.585). Pairwise Wilcoxon rank-sum tests indicated that cross-layer dissimilarity was significantly different among all three groups ($p < 0.001$).

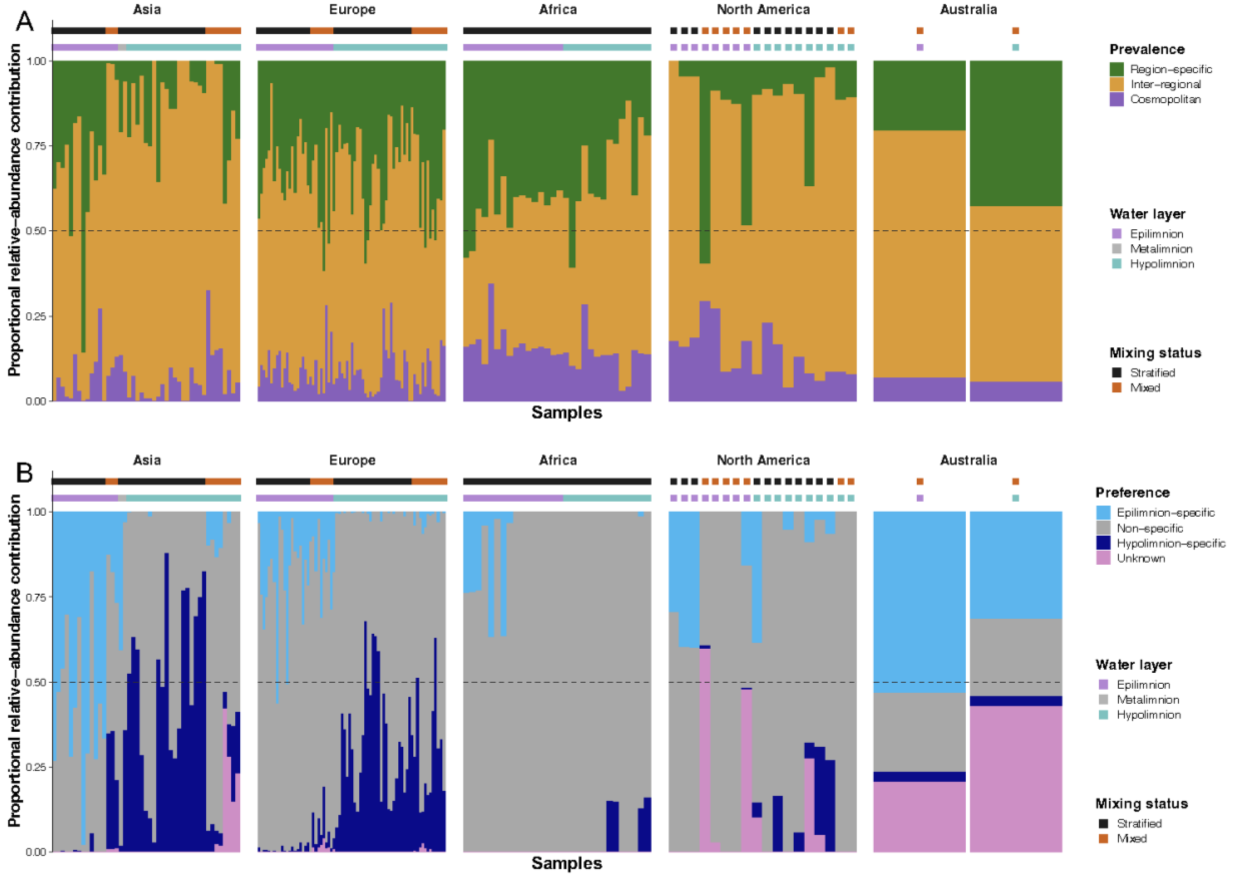


**Supplementary Figure 6. Relative abundance contribution of GV categories across samples.** Each stacked bar represents an individual sample, with bar heights normalized to the proportional abundance contribution of metagenome-assembled viral populations within each sample. Upper annotation rows indicate mixing status and water layer. The dashed horizontal line in both panels indicates the 50% proportional contribution threshold. (A) Relative abundance contribution of prevalence categories, including region-specific, inter-regional, and cosmopolitan viral populations. Cosmopolitan and inter-regional viruses combined contributed the largest fraction across samples. Inter-regional viral populations contributed variably among continents. (B) Relative abundance contribution of habitat preference categories, including epilimnion-specific, non-specific, hypolimnion-specific, and unknown viral populations. Epilimnion-specific viruses dominated most samples, whereas hypolimnion-specific viruses showed stronger contributions in deeper waters and in some high-latitude or stratified systems.

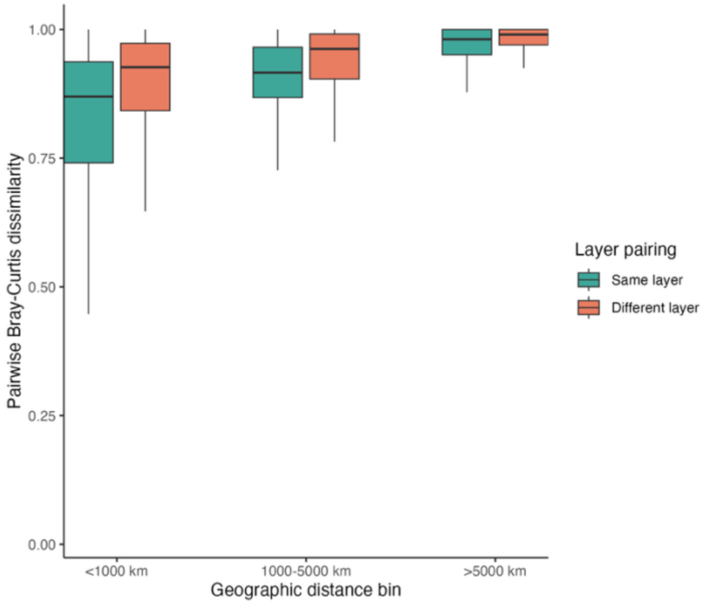


**Supplementary Figure 7. Pairwise Bray-Curtis dissimilarity across geographic distance scales and water layer pairings.** Boxplots show pairwise Bray-Curtis dissimilarities among stratified epi- and hypolimnion samples grouped by geographic distance bins (<1000 km, 1000-5000 km, and >5000 km). Comparisons are separated by layer pairing type: samples from the same layer (epilimnion-epilimnion or hypolimnion-hypolimnion) and different layers (epilimnion-hypolimnion). Bray-Curtis dissimilarity generally increased with geographic distance, becoming consistently high at intercontinental scales (>5000 km). Opposite-layer comparisons consistently exhibited slightly higher dissimilarity than same-layer comparisons across all distance bins.

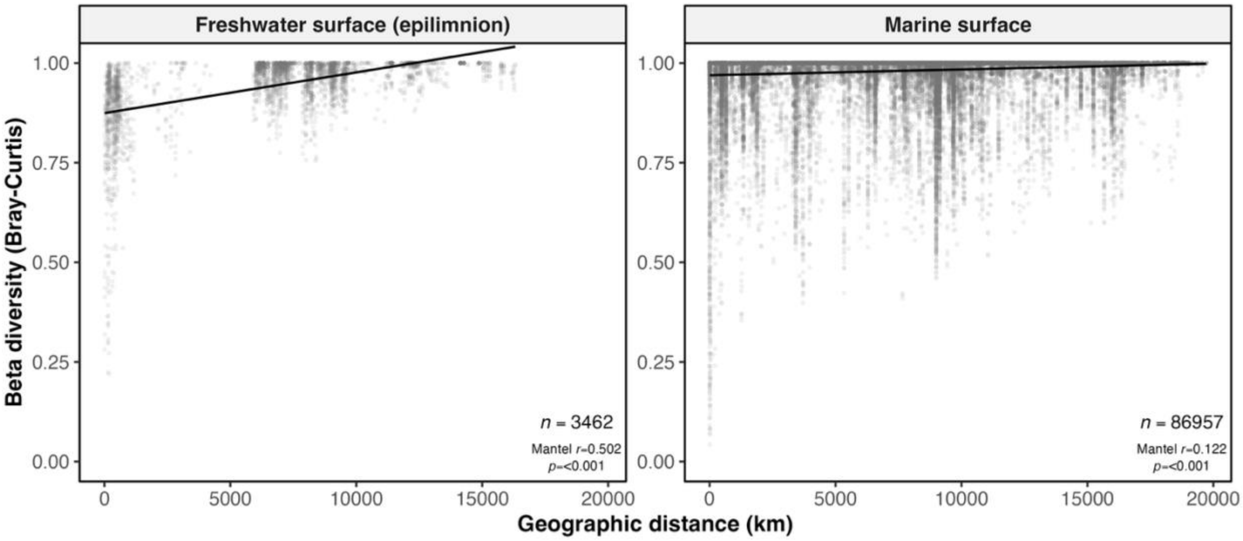


**Supplementary Figure 8. Distance-decay relationships of GV communities in freshwater and marine surface environments.** Beta diversity was measured using Bray-Curtis dissimilarity, and its relationship with geographic distance was assessed using Spearman's rank correlation. Spearman's rank correlations assessed through Mantel tests are given on the lower right of each plot. Linear regressions are shown as black lines. These regressions were not used to infer correlations between community dissimilarity and geographic distance but such correlations are quantified by the Mantel test results.

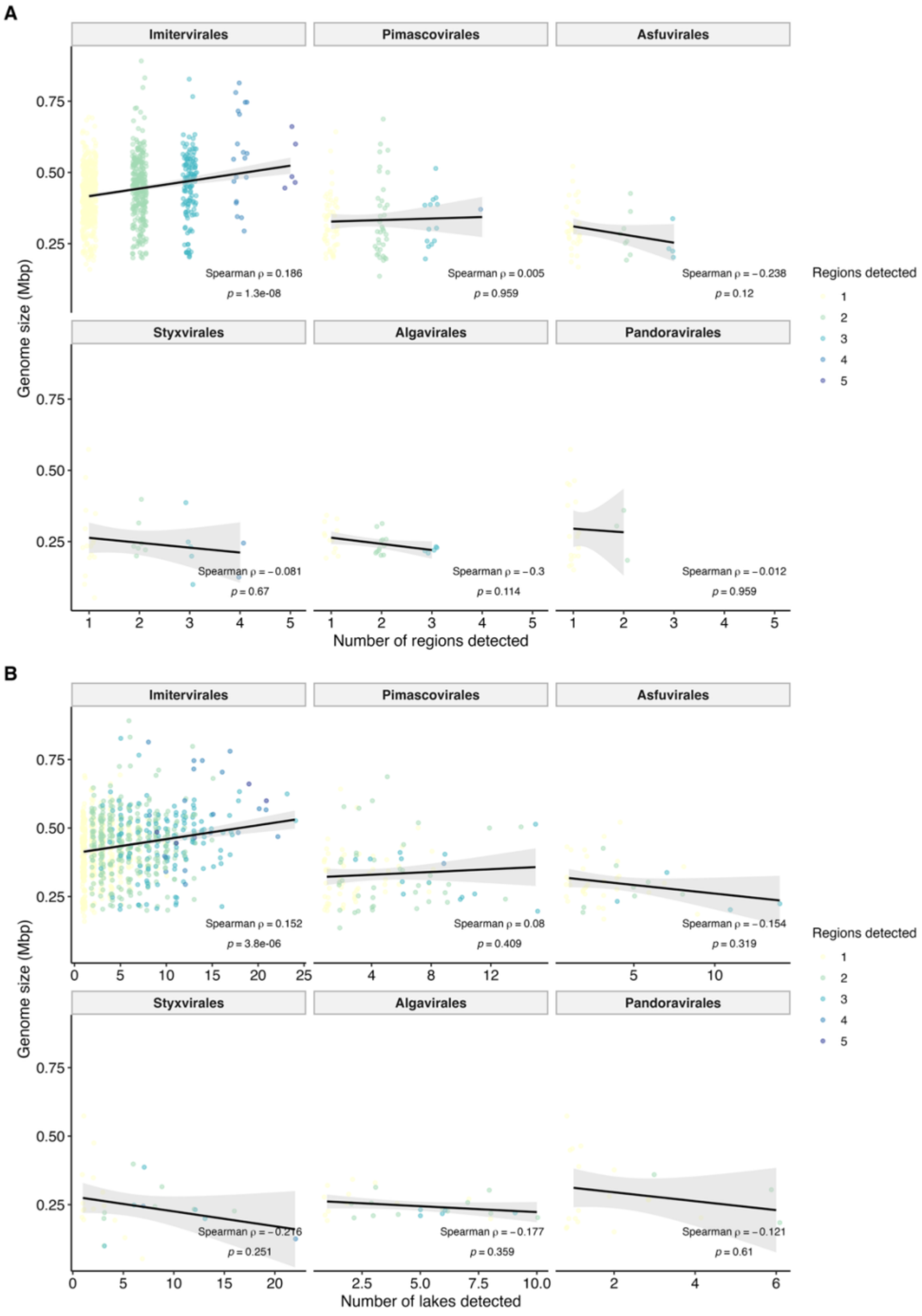


**Supplementary Figure 9. Relationship between giant virus genome size and geographic occupancy across viral orders.** (A) and (B) shows correlations between genome size and the number of regions and the number of lakes in which each GV MAG was detected, respectively. Each point represents a GV MAG and is colored according to number of continents detected in panel B. Black lines indicate linear regression fits with 95% confidence intervals shown in gray. Spearman's rank correlation coefficients (ρ) and associated p-values are shown within each panel. A significant positive relationship between genome size and geographic occupancy was observed only within *Imitervirales*, whereas other viral orders showed weak or nonsignificant associations.

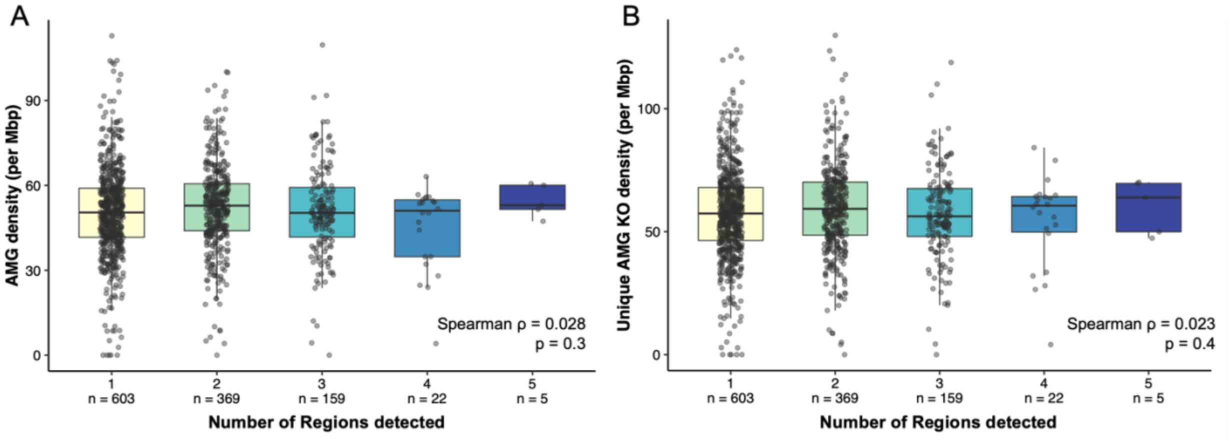


**Supplementary Figure 10. AMG density by number of detected regions.** Boxplots with overlaid individual data points show (A) counted AMG density and (B) unique AMG KOs density per Mbp across samples grouped by the number of regions detected. Sample sizes are shown below each group. Spearman correlation tests indicate no significant association between region number and AMG density (counted AMG density, $\rho = 0.028$, $p = 0.3$; unique AMG KO density, $\rho = 0.023$, $p = 0.4$).

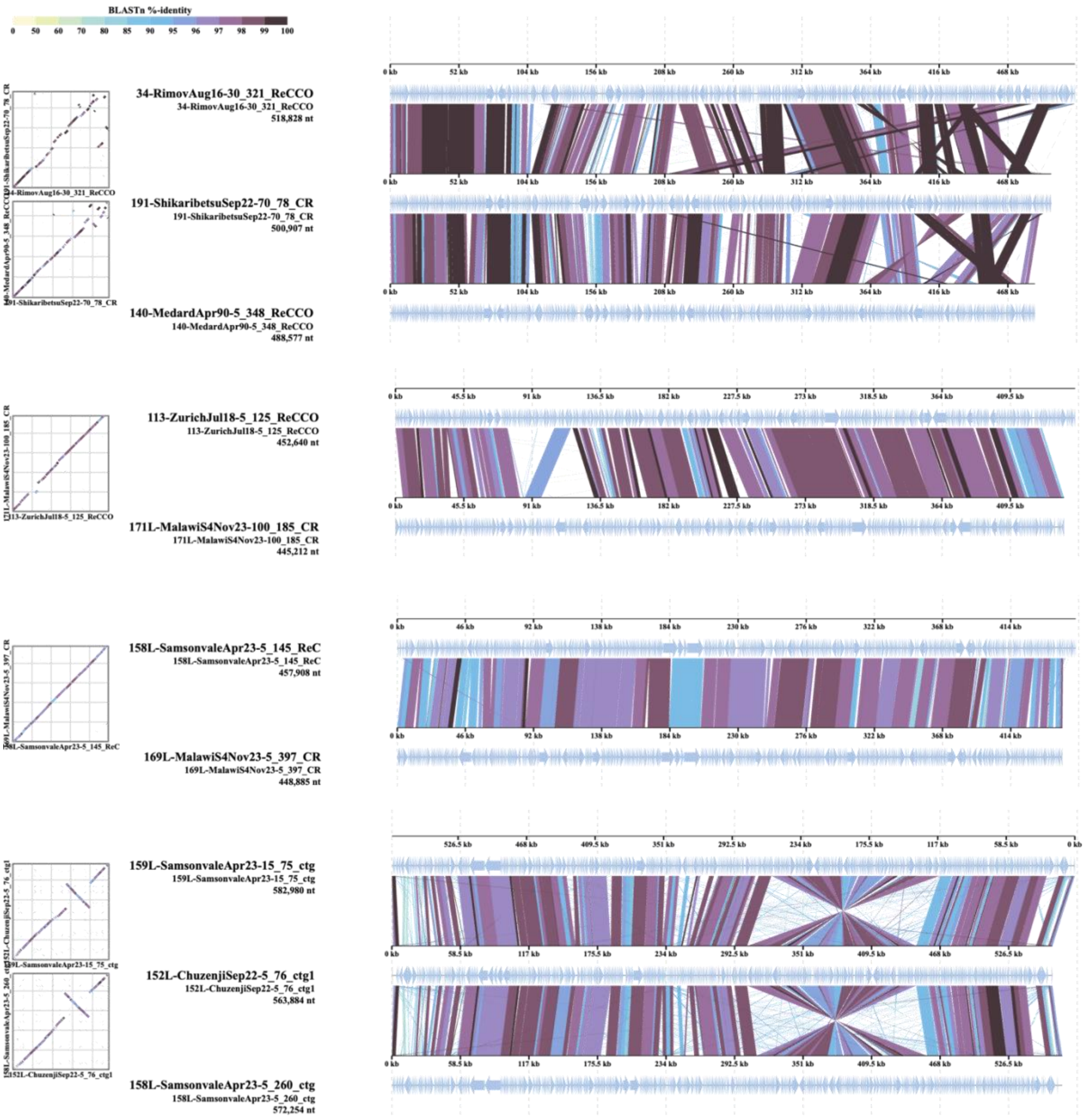


**Supplementary Figure 11. Genome alignments of GV MAGs recovered across continents colored by BLASTn identity percentages.** Genome pairs with ANI > 97% and aligned fraction > 80% were selected for plotting.

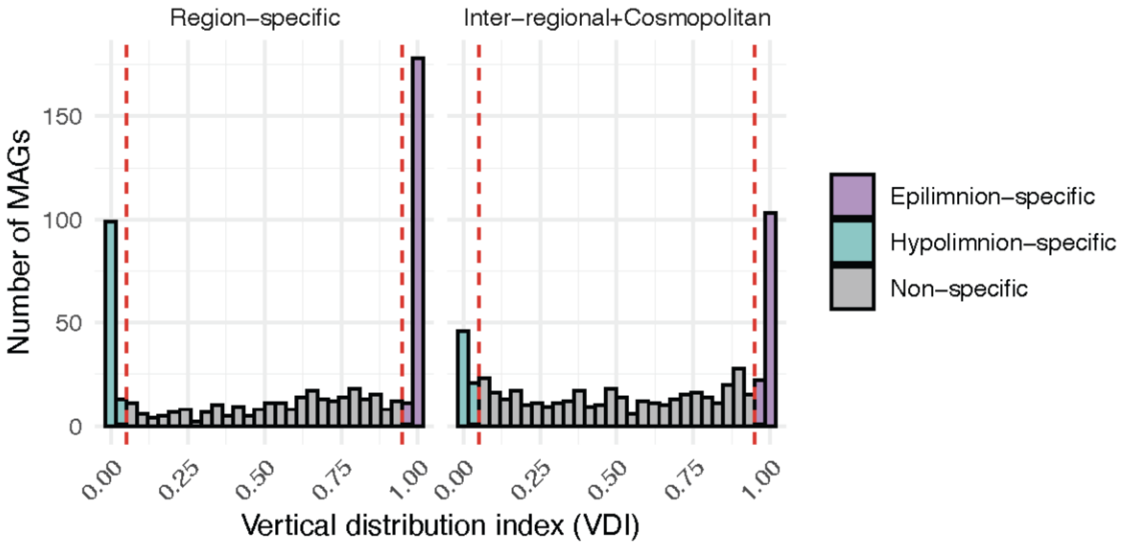


**Supplementary Figure 12. Distribution of vertical habitat preference across GV MAGs with different geographic breadths.** Histograms show the distribution of the vertical distribution index (VDI) for region-specific MAGs (detected in only one continent) and inter-regional/cosmopolitan MAGs (detected in >1 continents). VDI was calculated as the proportion of total abundance contributed by epilimnion samples across all stratified samples (see Methods). MAGs with VDI > 0.95 were classified as epilimnion-specific, whereas MAGs with VDI < 0.05 were classified as hypolimnion-specific. Intermediate values indicate non-specific MAGs. Histogram bars are colored according to vertical habitat preference categories. Red dashed lines indicate the classification thresholds at VDI = 0.05 and 0.95.

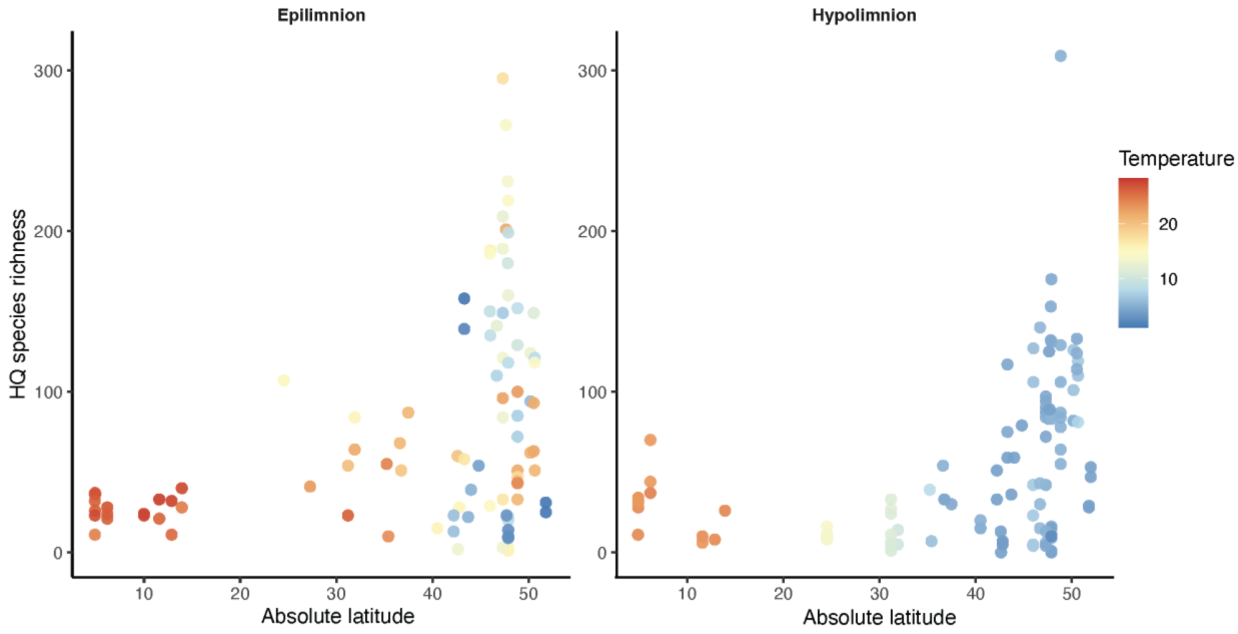


**Supplementary Figure 13. Relationship between absolute latitude and species richness across two lake water layers: the epilimnion (left) and hypolimnion (right).** Each point represents an individual sampling site. Color indicates water temperature (°C).

### Supplementary Tables and Legends

**Supplementary Table 1. Description of the 203 metagenomic samples.** This table summarizes the 203 metagenomic datasets included in this study, including sampling lake, sampling site, lake origin, sampling date, water layer, continent, country, sampling depth, lake trophic status, latitude and longitude, water stratification status, water temperature, data source, and sequencing read information.

**Supplementary Table 2. Statistics of the 1,663 GV MAGs recovered in this study.** For each MAG, this table reports the number of contigs, genome size, N50, GC content, taxonomic order, taxonomic family, completeness estimated by GVclass, quality classification, number of detected samples, number of detected lakes, number of detected continents, geographic prevalence classification, and vertical habitat preference.

**Supplementary Table 3. Terminal inverted repeat (TIR) information.** TIR length, coordinates, and sequence identity are reported for each detected TIR.

**Supplementary Table 4. Average nucleotide identity (ANI) values for nucleocytovirus MAGs.** Pairwise ANI comparisons were performed between each nucleocytovirus MAG and reference GV genomes in the GV MAG v2 database (see Methods). Only the highest-ANI match for each MAG is reported. The source ecosystem of each reference genome is also provided. Genome pairs with ANI >95% are highlighted. MAGs without a detectable ANI match because of high sequence divergence from the reference database are not included.

**Supplementary Table 5. ANI values for mirusvirus MAGs.** Pairwise ANI comparisons were performed between each mirusvirus MAG and reference GV genomes in a public database (see Methods). Only the highest-ANI match for each MAG is reported. The source ecosystem of each reference genome is also provided. Genome pairs with ANI >95% are highlighted. MAGs without a detectable ANI match because of high sequence divergence from the reference database are not included.

**Supplementary Table 6. Abundance profiles of the 33 mirusvirus and 1,126 nucleocytovirus MAGs.** Reads per kilobase of genome per million mapped reads (RPKM) are reported for each GV MAG across all 203 metagenomic samples.

**Supplementary Table 7. Orthogroups specific to inter-regional or cosmopolitan GV MAGs.** This table lists orthogroups detected exclusively in inter-regional or cosmopolitan GV MAGs. Functional annotations generated by eggNOG-mapper and the number of MAGs containing each orthogroup are provided.

**Supplementary Table 8. ANI values for all 1,630 nucleocytovirus and 33 mirusvirus MAGs.** Pairwise ANI comparisons were performed between each GV MAG and the Lake Biwa giant virus MAG (LBGVMAG) dataset (see Methods). Only the highest-ANI match for each MAG is reported, and genome pairs with ANI >95% are highlighted. Comparisons of vertical habitat preferences between this study and the Lake Biwa dataset are also included. MAGs without a detectable ANI match because of high sequence divergence are not included.

**Supplementary Table 9. Species richness of mirusviruses and high-quality nucleocytoviruses in each sample.** This table reports the number of detected mirusvirus and high-quality nucleocytovirus species (represented by dereplicated MAGs) in each metagenomic sample and was used to generate Fig. 3C and Fig. S13. The sampling month and season of each sample are also provided.

### Supplementary Methods

#### Decontamination of MAGs

For nucleocytovirus MAGs, contigs without GV signature were removed using a weighted confidence scoring system. Each contig was annotated independently with VirSorter2 v2.2.4 (--include-groups NCLDV, dsDNAphage --min-length 2500 --min-score 0.5) [1], the contig mode of ViralRecall v2.1 (-db GVOG) [2], and the end-to-end pipeline of geNomad v1.8.0 [3]. A confidence score was assigned to each contig based on the following criteria: a score of +1 was added if any of the three tools classified the contig as a GV, while −1 was applied if VirSorter2 or geNomad annotated it as a bacteriophage. Detection of any of the seven conserved nucleocytovirus marker genes (*Jelly-roll MCP, PolB, SFIIB, A32, VLTF3, TopII,* and *RNAPL*) contributed an additional +1 to the score. Furthermore, a penalty of −0.5 was imposed if CheckV v1.0.1 [4] detected no viral-specific genes but at least one cellular-specific gene on the contig. Contigs with a final confidence score below two were regarded as non-GV and excluded from the MAG in this step.

For mirusviruses, we employed public HMM profiles constructed from a comprehensive database of mirusvirus orthologous groups [5]. Hmmsearch [6] was used to screen all contigs against this database, and only contigs with at least one significant hit were retained (e-value = $1 \times 10^{-5}$).

#### Quality assessment

MAGs were classified as high-quality if they met the following criteria: completeness >70%, GVOG8df <2, order-level duplication (order_dup) <1.5, number of contigs <30, and N50 >5000 bp [7]. Medium-quality MAGs were also retained if they satisfied completeness >30%, GVOG8df <3, order_dup <2, number of contigs <50, and N50 >5000 bp. To assess linear complete genomes, we identified putative terminal inverted repeats through inverted alignment by Minimap2 v2.26 [13]. We then filtered the alignment result with minimum length of 99 bp and minimum sequence identity of 0.9. Only if the inverted sequences were aligned perfectly from at least one end of the genome and the length of the other flanking end was shorter than the length of aligned sequence, we considered it a terminal inverted repeat. For circularity assessment, we selected single-contig MAGs. Direct repeats (DIRs) were detected using CheckV v1.0.1 [4] for short-read-assembled MAGs, whereas circularity of long-read-assembled MAGs was evaluated based on the Flye assembly output [8].

#### Novelty assessment

To assess the novelty of this curated database, we clustered the resulted non-redundant high- and medium-quality GV MAGs with previously reported GV MAG datasets (Giant Virus MAGs v2 [9] and LBGVMAGs [10]) using FastANI v1.33 [11].

#### Orthogroup reconstruction

To identify auxiliary metabolic genes (AMGs), we clustered protein sequences predicted from the MAGs into orthogroups (OGs) using OrthoFinder v3.0.1b1 [12]. The longest sequence from each OG was selected as the representative and annotated with KEGG Orthologs (KOs) [13] using eggNOG-mapper v2.1.13[14].

### Phylogenetic inference

A concatenated alignment of seven marker genes (*PolB, TFIIB, TopoII, A32, SFII, VLTF3,* and *RNAPL*) was generated from the dereplicated nucleocytovirus MAGs and isolated nucleocytovirus reference genomes using "ncldv_markersearch" script [15]. Alignments were trimmed using trimAl v1.5.0 (90% gap threshold) [16] and phylogenetic trees were inferred using IQ-TREE v3.0.1 (-m MFP -B 1000 -alrt 1000) [17]. The best-fitting model (LG+F+I+R10) was selected according to the Bayesian information criterion from the ModelFinder [18]. For mirusviruses, we included a recently curated database [19] for reference and built the tree using HK97 MCP gene. Alignment trimming and tree building were done in the same manner as nucleocytoviruses and LG+F+R10 model was selected to be the best-fitting model according to the Bayesian information criterion from the ModelFinder [18].

### PERMANOVA test on the Bray–Curtis dissimilarity matrices

Community compositional differences between African and non-African samples and between water layers were tested using permutational multivariate analysis of variance (PERMANOVA) implemented in the adonis2 function of the vegan package [20, 21] on Bray–Curtis dissimilarity matrices, with significance assessed using 999 permutations.
